# Design of overlapping genes using deep generative models of protein sequences

**DOI:** 10.1101/2025.05.06.652464

**Authors:** Gun Woo Byeon, Marc Expòsit, David Baker, Georg Seelig

## Abstract

In nature, viruses frequently evolve overlapping genes (OLG) in alternate reading frames of the same nucleotide sequence despite the drastically reduced protein sequence space resulting from the sharing of codon nucleotides. Their existence leads one to wonder whether amino acid sequences are sufficiently degenerate with respect to protein folding to broadly allow arbitrary pairs of functional proteins to be overlapped. Here, we investigate this question by engineering synthetic OLGs using state-of-the-art generative models. To evaluate the approach, we first design overlapped sequences targeting two different protein families. We then encode distinct highly ordered de novo protein structures and observe surprisingly high in silico and experimental success rates. This demonstrates that the overlap constraints under the structure of the standard genetic code do not significantly restrict simultaneous accommodation of well defined 3D folds in alternative reading frames. Our work suggests that OLG sequences may be frequently accessible in nature and could be readily exploited to compress and constrain synthetic genetic circuits.

## Introduction

In 1977, Frederick Sanger and colleagues sequenced a whole genome for the first time in history - the 5.4kb DNA of ΦX174 bacteriophage^1^. This result resolved a head-scratching problem that had been bothering scientists for some time where the total summed length of the proteins produced by ΦX174 was too much to fit into the measured size of its DNA. Previous measurements had not been wrong, but “pairs of genes are coded by the same region of DNA using different reading frames”. Fast-forwarding to the current era of genomics, overlapping genes (OLG) have now become a familiar phenomenon especially in virology, where more than half of all known viruses encode at least one OLG in their genomes^2^. Within individual genomes, the extent of overlap can be quite large. In the well-established example of hepatitis B virus, every single known gene overlaps at least one other gene, accounting for ∼50% of its 3.2kb genome^3^. Outside of viruses, improved mass spectrometry and ribosome profiling methods have rapidly increased the number of candidate OLGs across a wide variety of cellular phylogenies, and it is now clear that many alternative open reading frames currently missing in the reference gene annotation databases are expressed and functional^4,5^.

These observations altogether suggest that OLGs arise in natural genomes at a surprisingly high frequency in contrast to the naive intuition that coding two proteins on the same DNA must be severely restrictive since each nucleotide is part of two codons. One explanation for this paradox has been that OLGs have a tendency to encode proteins with higher intrinsic structural disorder (ISD)^6–8^. Since sequences with ISD are more degenerate and thus more tolerant of mutations than sequences with well defined 3D structures, an ISD-rich protein in one frame would alleviate the evolutionary constraint on the other. On the other hand, native protein structures have been successfully encoded by a reduced amino acid alphabet, using as few as five unique amino acids^9^. This suggests that the minimal sequence complexity required to fold unique structures is significantly less than the full complexity that can be achieved with 20 amino acids, which raises the possibility that OLG sequences encoding highly ordered structures are readily feasible biophysically even if their evolutionary landscape may be more rugged. Is the sequence space of protein folding sufficiently degenerate to allow encoding arbitrary pairs of functional proteins in overlapping reading frames? How difficult is it for evolution to sample OLGs in nature? These questions motivate a synthetic biology approach to designing OLGs from scratch.

Synthetic OLGs also have obvious potential for biotechnology applications. For example, the ability to rewrite genes to overlap each other will aid compact genome engineering by reducing the footprint of genes and payloads. Furthermore, overlapping a gene of interest - such as a key catalytic enzyme in a metabolic engineering project - with an essential selection gene can ensure security of synthetic gene circuits by increasing genetic stability and/or introducing containment measures^10^.

The earliest attempt at synthetic OLGs used Rosetta to encode the active sites of Class I and II aminoacyl-tRNA synthetases on opposing strands with reverse complement codons, with the goal of stabilizing protein backbones around existing overlap-compatible ATP-binding peptide motifs that are the basis of Rodin-Ohno dual-coding origin hypothesis for tRNA synthetases^11,12^. A more generalizable algorithm was then developed by Opuu et al., which generates synthetic overlapping homologs by maximizing BLOSUM similarity scores to the target proteins^13^. Most recently, CAMEOS algorithm was developed to incorporate direct-coupling analysis models of protein sequences and was successful at designing a number of synthetic OLGs with genetic biocontainment applications^14^.

In recent years, deep learning (DL) methods have facilitated large advances in modeling of protein sequences, leading the DL models to become the top performers in various prediction or *de novo* design tasks^15,16^. We wondered whether we could exploit DL models to improve engineering of synthetic OLGs. Here, we describe a computational algorithm that enables the use of current state-of-art generative models of protein sequences for OLG design. We first illustrate a design scenario targeting a pair of biosynthetic and essential genes for entanglement conditioned on homologous protein families. We show that the resultant synthetic sequences are highly diverse and divergent from natural sequences yet score as well as them on a number of *in silico* benchmarks. Secondly, we generate OLG sequences encoding pairs of highly ordered structures, conditioning on their backbone atom coordinates. We find that the qualities of overlapping vs. non-overlapping designed sequences are mostly on par *in silico*, a striking observation further bolstered by high experimental success rates in their validation by recombinant expression and structural characterization. These results counter the notion that well defined 3D protein folds are rarely overlap-encodable in alternative reading frames without segregation, providing evidence for high biophysical and evolutionary feasibility of OLGs under the standard genetic code.

## Results

### A computational algorithm for designing OLG encoding sequences

We are designing two protein sequences simultaneously, and we want to jointly maximize the fitness or likelihood scores of each sequence. However, the fundamental challenge of OLG design is that because of the interdependence of the codons in the alternative frames, only a tiny fraction of the full sequence space is actually available as potential solutions. Note that there are 5 alternative reading frames given a nucleotide sequence and each with different constraints: 2 plus-strand and 3 minus-strand, denoted +1, +2, -0, -1 and -2 frames with +0 as the “reference” frame (**Fig. 1A**).

**Figure 1.**
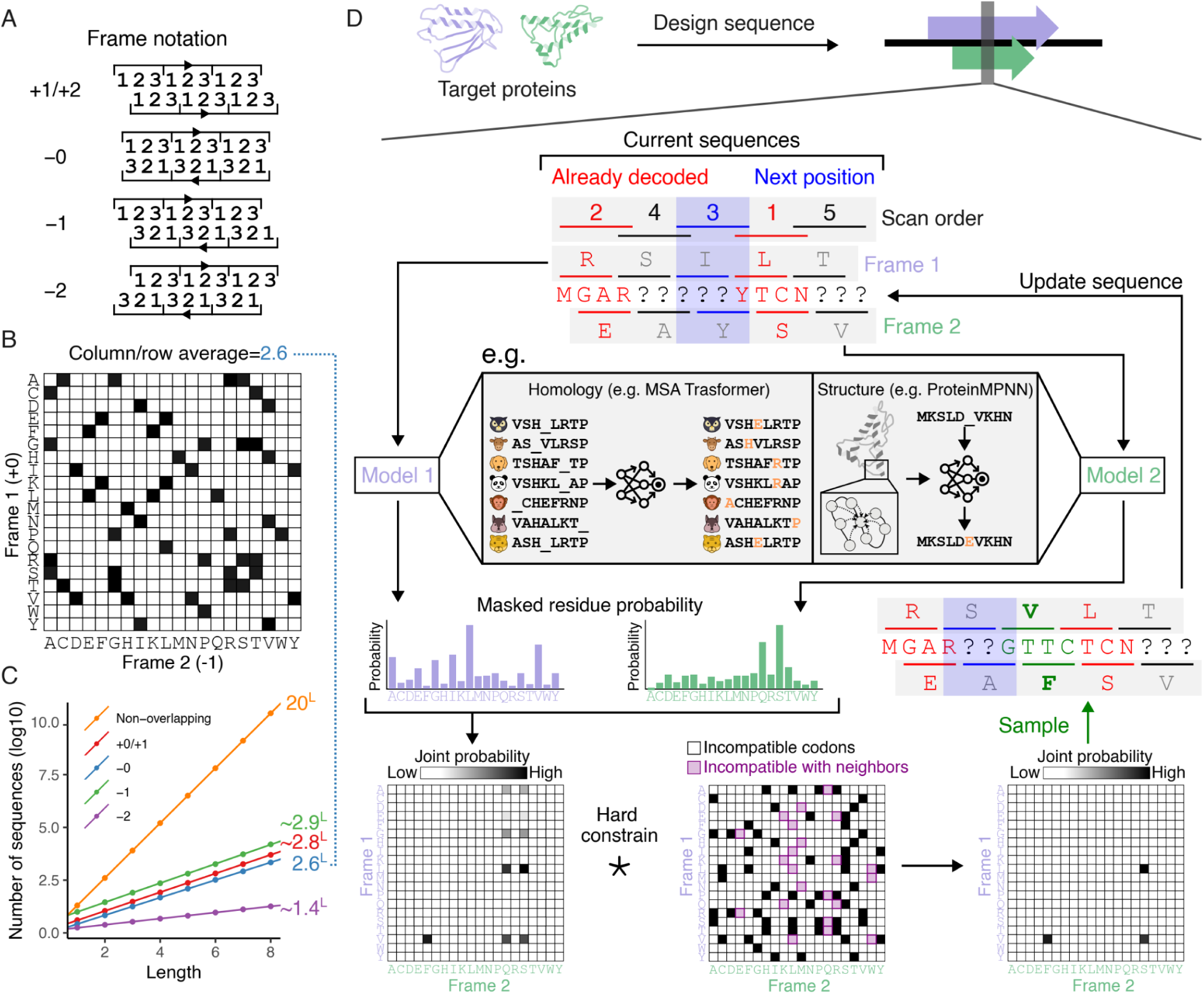
Computational design of synthetic overlapping genes. (**A**) The 5 possible alternative reading frames for encoding a pair of proteins in the same nucleotide sequence, which reduces to 4 possible unique types of constraints due to reciprocity in the two plus strand frames - for example, if sequence S’ is +1 relative to sequence S, then S is +2 relative to S’. (**B**) A two-dimensional matrix illustrating pairs of amino acids that are compatible in the -0 alternative frame arrangement. Filled cells indicate that there is at least one trinucleotide that codes for the pair of amino acids indicated in the rows and columns in the opposite strands. (**C**) Reduced sequence space of overlapping genes. Given a random sequence S with specified lengths on the x-axis, all possible sequences uninterrupted by stop codon in each alternative reading frame are enumerated, and the average number of total possibilities from N=2000 are plotted on the y-axis. (**D**) A schematic illustrating the constrained iterative sampling algorithm for designing overlapping genes using discrete masked models of protein sequences. The nucleotide sequence is represented as a series of linked quartets rather than triplets to fully describe phase-shifted alternative reading frames position-wise. One position is sampled per iteration for each protein, where the two positions are the residues translated by the overlapping triplet codons. At each step, a forward pass on the generative model (for example, MSA Transformer for protein families or ProteinMPNN for structures) gives the amino acid probability vector for the current position of each target protein. The pairwise product of the two amino acid probability vectors returned by the respective models gives the joint probability. Incompatible pairs of amino acids according to the frame arrangement and the codon table are hard constrained by masking them. When the amino acid pair in the neighboring position(s) have already been chosen, the joint probability matrix is additionally masked at the pairs that are incompatible with the neighboring residues. The amino acid pair is sampled from this hard-constrained joint probability matrix, and the respective sequences are updated by replacing the current tokens with the sampled pair of tokens.

To first give an intuition into the constraint on the overlap and illustrate its magnitude, we analyze the -0 frame. Quantitative description of the -0 frame is relatively simple since it does not have a phase shift: the constraints at any position are independent of its neighboring positions. At each position, the choice of an amino acid in the reference frame corresponds to a set of compatible amino acids in the alternative frame according to the codon table, which can be visualized as a two dimensional matrix of pairwise compatibilities (**Fig. 1B**). This shows that given a sequence in the reference frame, there are 2.6 possible choices per position in the alternative frame on average and thus 2.6^length^ possible sequences. It also follows that there are 2.6x20=52 compatible pairs of amino acids out of 20x20=400 combinations that are available at each position which means there are 52^length^ possible overlapping pairs of sequences. These are minuscule fractions of the full space of all 20^length^ sequences or 400^length^ pairs of sequences but nevertheless represents a large space where specifying one sequence does not fully specify the other due to the degeneracy of the genetic code. Other alternative frames are non-trivial to analyze but we can derive analogous measures of degeneracy using Monte Carlo approximation. We find that +1 and -1 frames have more degrees of freedom than -0 frame (∼2.8 and ∼2.9 respectively) while -2 frame is significantly worse (∼1.4) compared to the other frames (**Fig. 1C**).

Similar to how the overlap constraints can be decomposed position-wise, we can also simplify the problem of sampling overlapping protein sequences from generative models by using a position-by-position sampling procedure. Currently, many top-performing generative models of protein sequences are trained with discrete token masking objectives and learn the conditional distribution of masked tokens given partially masked sequence^17^. Designing new sequences with such models uses an iterative decoding scheme. We therefore reasoned that we can iteratively sample the two protein sequences simultaneously from the respective model instances while ensuring the compatibility of the amino acid pairs in alternative reading frames which can be described position-wise as illustrated above.

The -0 frame is again the simplest here due to the lack of phase shift. At each step, an overlapping pair of positions are masked and predicted - one for each of the target proteins. The pairwise product of the two amino acid probability vectors at the masked overlapping positions gives a joint probability matrix, which is simply hard constrained by zeroing the incompatible amino acid pairs before sampling one. The trick lies in extending this strategy to other frames which do have a phase shift. To this end, we use a systematic scan such that each position is visited exactly once per scan and additionally constrain the joint probability by the amino acids in its neighboring positions, if the neighbors have been already sampled in the previous steps (**Fig. 1D**). This ensures that after a complete sweep through all positions in a number of steps equal to length, the pair of sequences are always overlap-compatible. In practice, we find that quality of the designs as scored by model pseudolikelihoods improves with multiple complete sweeps. Furthermore, biasing the scan order in successive sweeps by position-wise degeneracy of the amino acids or by position-wise probability of the chosen amino acid further improves the design quality (**Supplementary Fig. 1A-D**).

### Diversity and quality of OLG designs targeting natural protein families

To evaluate our method, we first consider a homology-conditioned design problem: can we design overlapping gene sequences that are highly probable members of target protein families? To demonstrate this capability, we choose a pair of bacterial genes as our overlap targets: chorismate mutase (CM) and translation initiation factor 1 (IF1) (**Fig. 2A**). This choice entangles a biosynthetic gene with an essential gene, representing a mutational restriction application scenario of synthetic OLGs ^14^. Both proteins are also similarly sized and structural knowledge is available to aid their *in silico* evaluation^18,19^.

**Figure 2.**
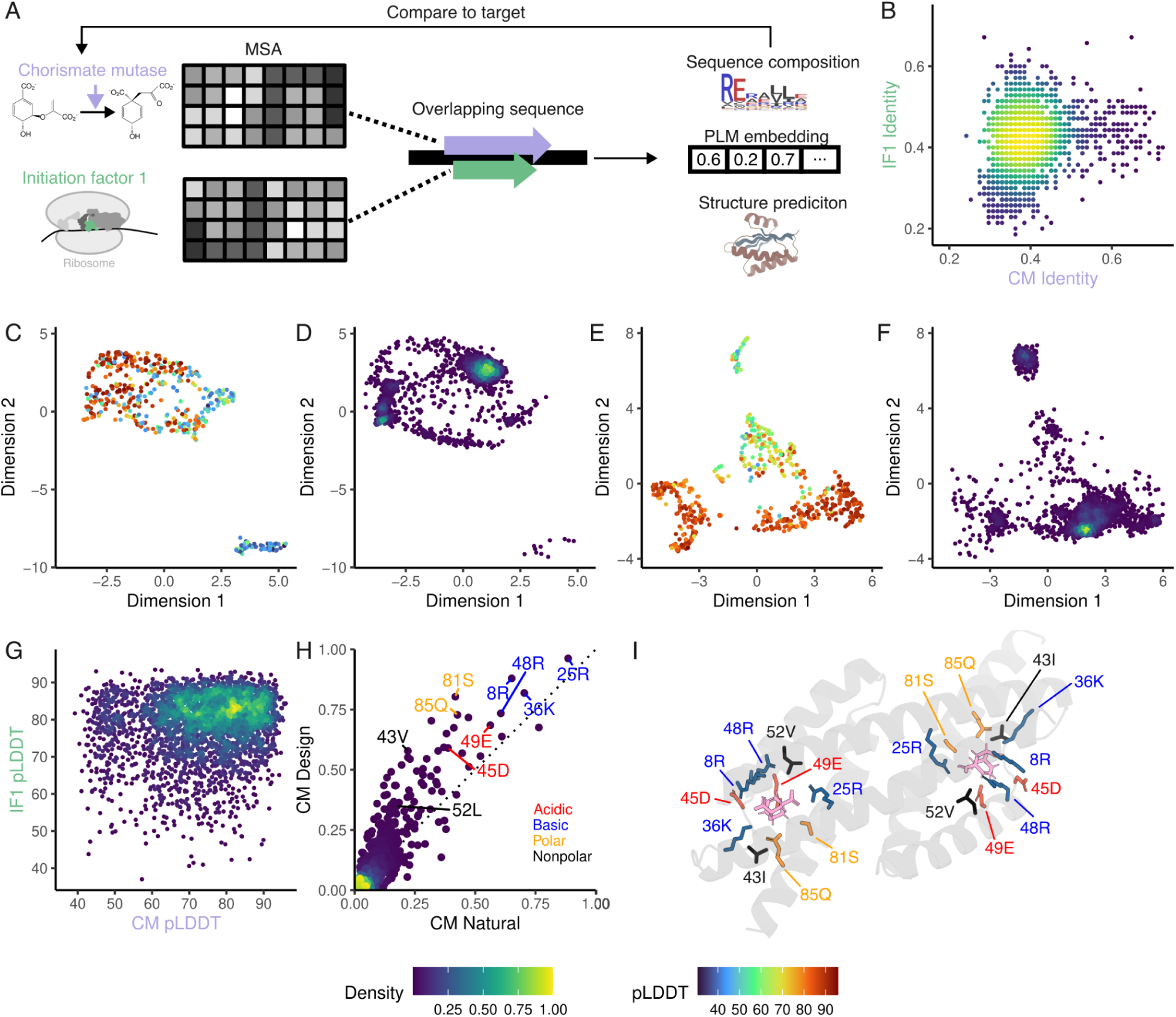
Targeting a pair of homologous protein families for OLG sequence design. (**A**) Target protein families - chorismate mutase (CM) and initiation factor 1 (IF1) - and the evaluation metrics. (**B**) A scatter plot of identities: each dot is a pair of OLG sequence designs, and on each axis are the identities to the closest natural sequences. Colored by density. (**C**) UMAP visualization of the ESM2 embeddings of the natural homologous sequences of CM. Colored by mean pLDDTs of the AlphaFold2 predictions for each sequence. (**D**) Projection of the ESM2 embeddings of the OLG synthetic homologs of CM in the same space of (C), colored by density. (**E**, **F**) Same as (C) and (D) but for IF1 sequences. (**G**) A scatter plot of AlphaFold2 pLDDTs of the structures predicted for the synthetic OLG sequences. Colored by density. (**H**) First-order statistics of the natural homologs of CM versus the designed CM sequences. Each dot is the proportion of the particular amino acid at a given position in the MSA of the sequences. Colored by acidic, basic, polar or nonpolar side chain types for the indicated active site residues within <3Å of the CM substrate. (**I**) The *E. coli* CM structure model (PDB: 1ECM) with the indicated active site residues in (H), which are identified as those <3Å in spatial distance to the substrate in the *E. coli* CM structure.

We use the iterative overlap-constrained sampling procedure described above with EvoDiff-MSA, which is a generative model with MSA Transformer architecture but trained using the order-agnostic autoregressive diffusion objective^20^. The MSA of each target protein is provided as a conditioning context, and the position-wise masking and constrained sampling procedure is performed on the first sequence of the input MSA. We generated 3307 fully overlapping sequence designs encompassing all possible offsets and alternative frame arrangements.

The designed sequences diverge significantly from natural sequences, with closest identities to natural sequences averaging at 38.9% and 42.3% for CM and IF1, respectively (**Fig. 2B**). We assessed whether these low identity designed sequences are credible members of the target protein families by analyzing their distribution in sequence space and comparing it to that of natural sequences in the same family. To this end, we embedded each generated and natural sequence using protein language models (ESM2, ESM3 and ProstT5)^21–23^. We visualized the embeddings of natural sequences in 2D via uniform manifold approximation and projection (UMAP) and projected the synthetic sequences as points onto that space (**Fig. 2C-F, Supplementary Fig. 2A-H**). We observed that the distributions closely align between the designed and natural sequences.

We next investigated whether the designed sequences form stable, expected folds by predicting their structures using AlphaFold2 (AF2) and computing the mean predicted local-distance difference test (pLDDT) scores, a measure of AF2’s confidence in its predictions ^24^. We specifically utilize the AF2Rank protocol, which executes AF2 predictions with a single sequence input along with a sidechain-masked structural template to evaluate the accuracy of the candidate structure provided as template^25^. We observe that designs in which both of the proteins in alternative frames exhibit high pLDDTs comparable to those of natural sequences (**Fig. 2G**). This suggests that the designed OLG sequences are structurally plausible members of the target protein families.

The embedding and structure analyses above in effect address the global resemblance of the designed proteins to natural proteins in the space of latent variables. We additionally evaluated the first-order, position-specific statistics of the designed sequences. This metric is especially pertinent to enzymes where the catalytic residues in active sites are often more strictly conserved for family members with the same reaction chemistry and substrate. We find that the position weight matrices (PWM) of designed protein sequences are highly similar to those of natural sequences, especially at the low entropy positions and in the case of CM, an enzyme, at the active site positions of the MSA (**Fig. 2H, I, Supplementary Fig. 2I-L**). These positions have a charged or polar amino acid as the most frequent residue and tend to have very low entropy in the PWM, likely comprising the catalytic residues critical for the reaction chemistry. This suggests that the designability of protein sequences under the constraints of overlapping codons are not broadly restrictive to retaining key residues required for functionality of the protein.

### Highly ordered protein backbone structures are easy to overlap

Encouraged by the high feasibility of OLGs given the naive initial expectations based on stringent overlap constraints, we set out to test the hypothesis that well defined 3D folds are readily overlapped. We compare the designability of overlapping and non-overlapping protein sequences for a set of highly ordered target structures (**Fig. 3A**). Specifically, we chose backbones generated by activation maximization of deep learning protein structure predictors^26^. These *de novo* structures represent energetically favorable, “ideal” versions of 3D folds existing in nature. We use the iterative overlap-constrained sampling procedure with ProteinMPNN, a well validated structure-conditioned generative model^27^. We generated 56,250 total overlapping sequence designs, which targeted all pairwise combinations of 15 backbones which are 100 amino acids long, spanning various α-only, β-only and mixed α-β fold classes, in all possible frame arrangements and all possible offsets at minimum overlap proportion of 95%. We also generated 33,000 non-overlapping designs for the same target folds at various sampling temperatures. We then asked whether AF2 predicts the designed sequences to fold to their respective target structures.

**Figure 3.**
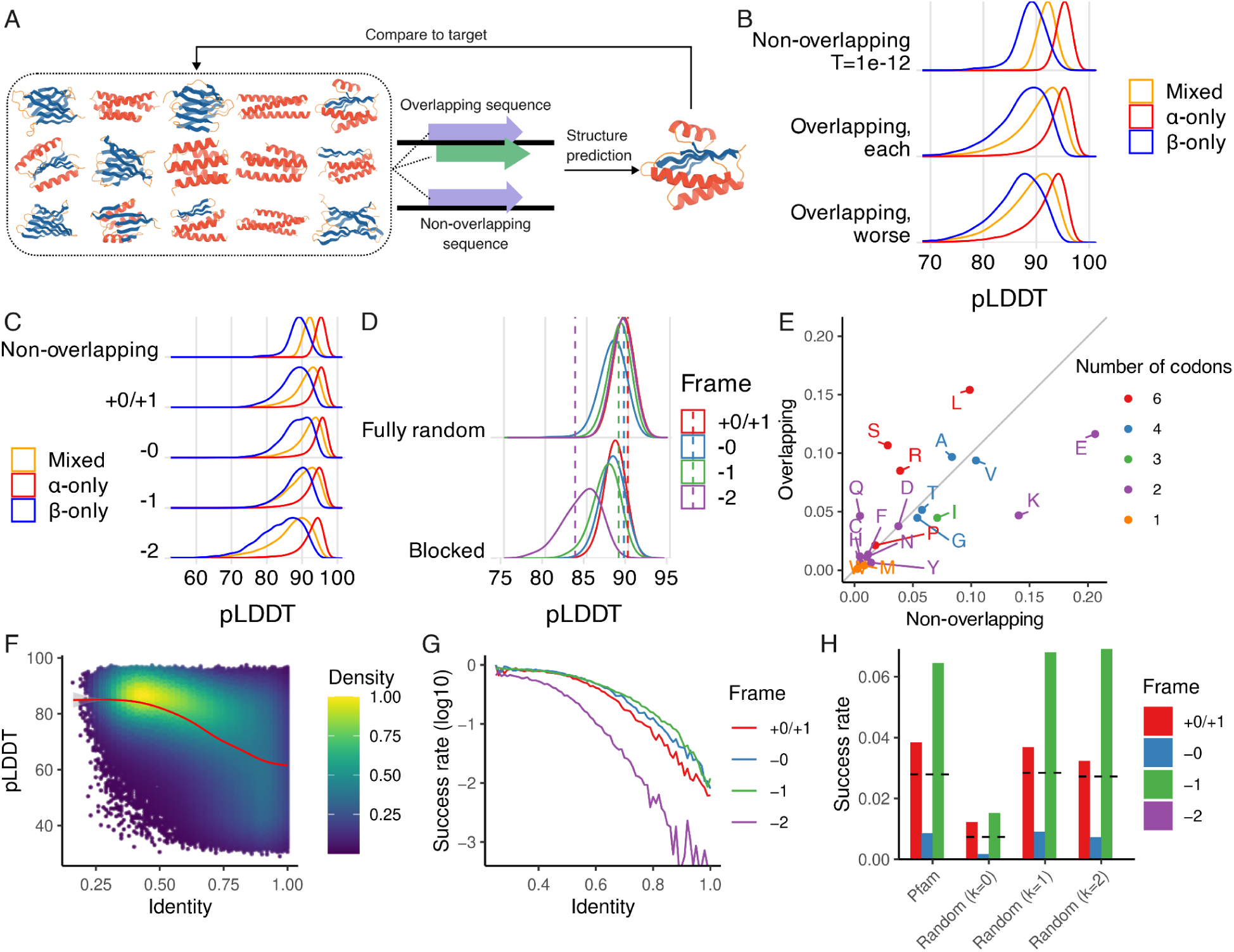
Designing OLG sequences encoding pairs of highly ordered protein structures. (**A**) Target backbone structures. (**B**) Distribution of pLDDTs for the AlphaFold2 predictions of ProteinMPNN designed sequences as overlapping pairs or standard non-overlapping proteins. Overlapping designs are shown altogether (middle), or only the worse pLDDTs of the overlapping pair are plotted (bottom). Colored by secondary structure classes. (**C**) Distribution of pLDDTs stratified by alternative frame arrangements. (**D**) Distribution of average pLDDTs for the AlphaFold2 predictions of overlapping sequence designs under randomized genetic codes (N=1000), either fully random (top) or maintaining block structure (bottom). Colored by alternative frame arrangements. Dashed lines indicate the average pLDDTs obtained from designs under the standard genetic code. (**E**) A scatter plot of the amino acid compositions of the designed overlapping (frame +0/+1) versus non-overlapping sequences. Each point labeled with amino acid letters and colored by the number of codons for the amino acid in the standard genetic code. (**F**) A scatter plot of the pLDDTs versus the specified identity of the parent protein in the reference frame to its seed sequence. Colored by density. (**G**) Same data in (F) plotted as success rate of designs above threshold (worse pLDDT >85 and TM score >0.7). Stratified and colored by alternative frame arrangements. (**H**) Proportion of success with natural protein domains from Pfam as the completely fixed parent protein in the reference frame or with random protein sequences that maintain zero-, first- or second-order compositional bias present in the set of Pfam domain sequences. Dashed lines indicate average across the frames.

We found that the pLDDTs for overlapping designs are nearly on par with the pLDDTs for non-overlapping designs - averaging at 90.2 versus 92.0 (**Fig. 3B**). Even when we examine only the worse pLDDT of each overlapping pair, the scores are only slightly lower - an average of 87.6. The similarity between the predicted and target structures, measured either by template modeling (TM) score or root mean square deviation (RMSD) are also on par between overlapping and non-overlapping designs (**Supplementary Fig. 3A, B**).

When examining these metrics stratified by the different alternative reading frames, we observed that -2 frame is particularly unfavorable (**Fig. 3C**). Intuitively, this may be explained by the block structure of the standard genetic code (SGC) which is mostly specified by the first and second nucleotides while the third position is largely degenerate. In -2 frame arrangement, the third position of one gene overlaps the third position of the other gene, which is an inefficient use of degeneracy and therefore likely leads to greater degree of incompatibility between the amino acid choices for the overlapping proteins. This is also in line with our initial estimation of the magnitude of overlap constraints as well as with previous formal analysis by others which have predicted -2 configuration to be the most restrictive^28–30^.

To investigate the impact of the structure of the SGC that may lead to such variations in the coding capacity of different overlapping reading frame configurations, we conducted the designability analysis using sets of randomized alternative genetic codes. We observed that fully randomized codon tables which do not maintain the block structure on average tend to be more favorable for encoding overlapping proteins and exhibit a bias against -0 configuration, whereas codes maintaining block structure typically exhibit significant bias against -2 configuration (**Fig. 3D**). Intriguingly, among the block structure code sets, we observe that the SGC is relatively optimal for encoding OLGs in all other configurations (8.5, 20.5 and 24 percentiles for +0/+1, -0 and -1 frames, respectively) at the expense of -2 frame (73 percentile).

The SGC also imparts a unique compositional bias on OLGs: amino acid residues with a high level of codon degeneracy in the SGC are preferred, substituting for amino acids with similar biochemical properties but with less codon degeneracy (**Fig. 3E**). The bias patterns significantly depend on the relative reading frame configurations (**Supplementary Fig. 3C-E**). Altogether, these analyses suggest that the redundancy in the SGC is sufficient to encode ordered protein backbone structures as OLGs and that it confers distinct biases in the feasibility and composition of OLG sequences in different reading frames.

### OLG sequences are evolutionarily accessible

Given these results, we sought to address the evolutionary accessibility of novel proteins in alternative reading frames of an already existing gene. We begin with a seed protein sequence in one of the reading frames, designed by low temperature sampling of ProteinMPNN for the set of various target structures above. We then design OLG protein sequences in the alternative frame while enforcing a range of sequence identities that must be maintained for the protein in the reference frame. This allows us to evaluate the difficulty of constructing OLGs at different thresholds of non-synonymous mutations to the parent protein.

Limiting the number of mutations to the parent protein decreases the probability of successfully designing OLGs against specified target structures (**Fig. 3F**). Surprisingly however, we find that even at very low numbers of mutations to the parent protein, we can still obtain many successful overlapping sequence designs. This is true even in the most extreme case where no mutation is allowed to the parent protein, where ∼1% of the trajectories result in pLDDT >85 with TM score >0.7 (**Fig. 3G**). Under the same criteria, at least one successful OLG sequence design was obtained for every parent protein. When we repeated the analysis with sequences of natural protein domains from Pfam as the completely fixed parent protein, we obtained ∼3% success rate (**Fig. 3H**). Notably, we obtain the same result when using random protein sequences that maintain only the first-order compositional bias present in the set of natural sequences. Altogether, these results show that proteins which are highly optimal in their structural stability or in their natural biological fitness can accommodate other proteins with well defined folds in alternative reading frames without requiring significant changes to their original sequences.

### High experimental validation rates of designed OLG sequences

Though sequences generated from the models we have used have demonstrated high experimental success rates in the context of non-overlapping design, the significant deviation in sequence composition for OLGs motivated us to experimentally validate the OLG designs and assess their failure modes^27,31^. To this end, we selected a subset of 192 overlapping sequences from the previous design campaign entangling a set of 15 *de novo* backbone structures for recombinant expression and structural characterization. We filtered against the poorest quality sequences with low pLDDT, high RMSD vs. target or higher spatial aggregation propensity; otherwise we made random selections stratified by structural classes (α-only, β-only, and mixed α-β) to obtain an equal representation (**Table S1**, **Supplementary Fig. 4A**).

We observed that 207/384 (54%) individual proteins (in either frame and across structural classes) expressed successfully (**Fig. 4A, B**). Three pairs belonging to different secondary structure groups (α-only/α-only, α-only/β-only, β-only/β-only) were selected for further characterization and observed to have circular dichroism profiles matching their expected secondary structure and to be highly thermostable, up to 95°C for most (**Fig. 4C**). The success rate varies by secondary structure content of the proteins, ranging from 77% in α helical proteins to 30% in proteins with only β sheets (**Fig. 4D**), in agreement with the observation that the de novo design of β sheets is more challenging^32^. Most of the designs were purified from the cytoplasm (36.5%), while some required refolding from inclusion bodies (17.4%), with an overall median soluble yield of 8.5mg/L of culture equivalent (**Fig. 4D, E**).

**Figure 4.**
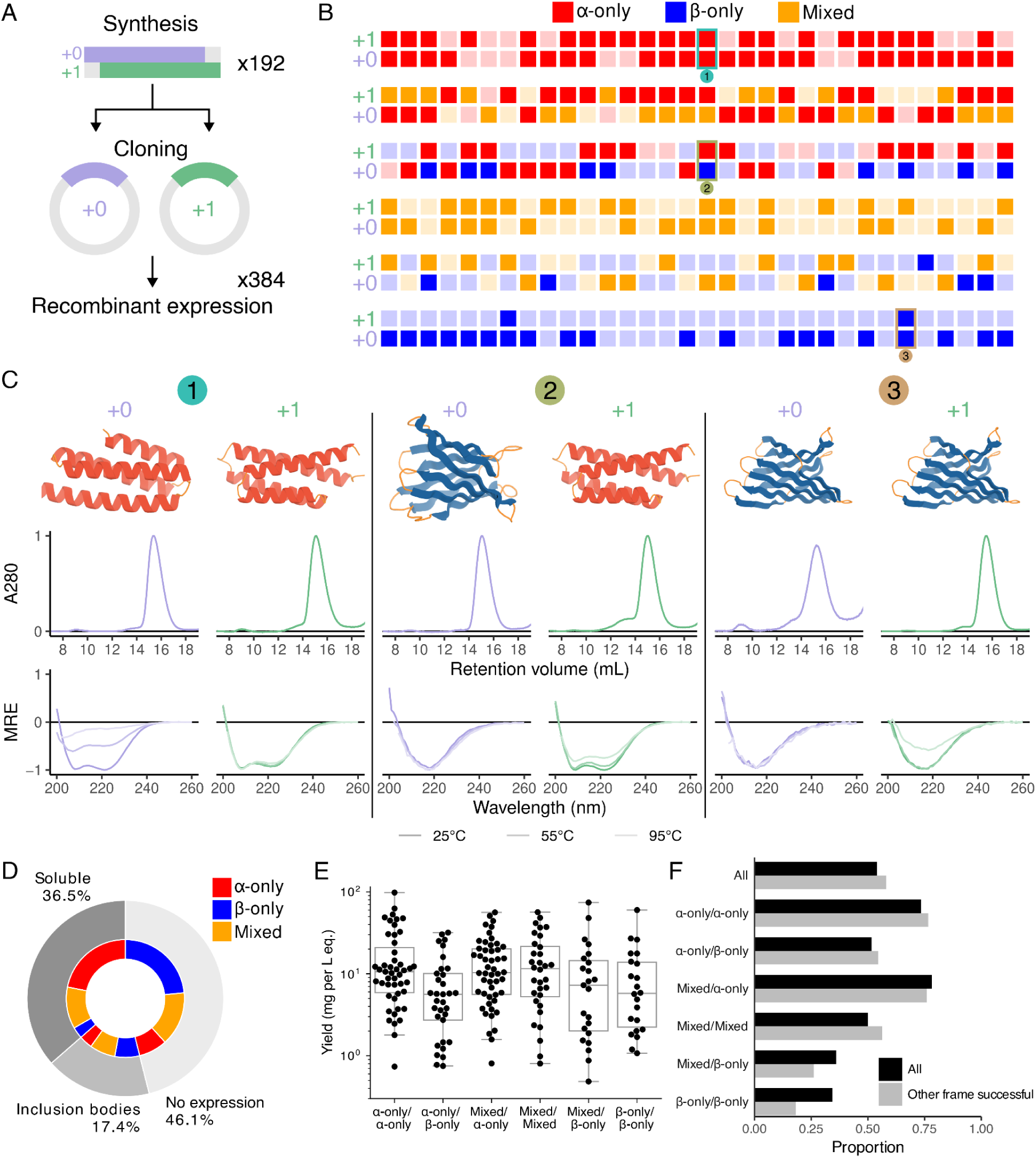
Experimental characterization of overlapping *de novo* protein pairs. (**A**) A schematic of the cloning - a total of 384 unique proteins are encoded as overlapping sequences over 192 DNA fragments, each which is cloned into two separate vectors which express the proteins in +0 or +1 frame. (**B**) Overview of successfully expressed proteins, grouped by secondary structure class and representing each pair next to each other vertically. The numbers indicate designed pairs selected for further characterization. (**C**) Biophysical analysis of representative overlapping protein pairs. Top: structures colored by secondary structure (red for α helix, blue for β sheets, and orange for loops). Middle: SEC traces after IMAC purification. Bottom: circular dichroism spectra at room temperature, 55°C or 95°C. (**D**) Percentage of designs that were successfully expressed from the cytoplasm, recovered from inclusion bodies, or not expressed. The interior pie chart indicates the secondary structure composition of proteins in each group. (**E**) Yield of purified protein of successful designs. (**F**) Percentage of successfully expressed designs by secondary structure classes considering individual designs (darker) or only those where the other frame expressed successfully (lighter).

When considering the success rate of the overlapping pairs, we observed 60/192 (31%) purified successfully, resulting in successful pairs across all possible combinations of secondary structure classes (**Fig. 4F**). Remarkably, we found that the success in one of the frames does not systematically impair success in the other frame, as the success rate when considering each protein individually is maintained when considering only those in which the other frame is successful. We also found that success is significantly associated with negative net charge of the proteins (**Supplementary Fig. 4B**). This suggests that the OLG-specific compositional bias induced by frequent use of high codon degeneracy amino acids likely impacted the solubility of recombinant proteins expressed in *E. coli*. Nevertheless, these results demonstrate that the restricted and distinctively biased sequence space of overlapping proteins does not severely compromise the high experimental validation rates observed for typical non-overlapping sequence designs with ProteinMPNN, further supporting our conclusion that OLG sequences encoding arbitrary pairs of well defined 3D folds are highly feasible.

## Discussion

Overlapping genes that encode separate proteins in two different reading frames of the same nucleotide sequence are surprisingly common in nature and found in a majority of viruses, but since their discovery in 1977, their very existence has posed a long standing biophysical and evolutionary puzzle. Here, we developed a computational algorithm that encodes two target proteins into the same DNA sequence under the constraints of the overlapping codons. Our study extends the prior efforts that used first- and second-order models and incorporates DL based generative modeling of protein sequences^13,14^. This allows us to look beyond templating the designs on existing natural protein families with large numbers of known homologous sequences. When we conditioned the designs explicitly on protein structures using an inverse folding model, we discovered that contrary to the common notions, stable protein backbones are readily dual encodable in alternative reading frames. Overall, we find that synthetic OLGs entangling arbitrary pairs of proteins are highly feasible to design through current protein sequence generation methodologies.

A key outstanding challenge for constructing synthetic OLGs will be to develop a robust approach to co-design DNA/RNA sequences alongside the overlapping protein sequences for OLG expression in the cell. Thus far, synthetic OLGs have only been expressed in prokaryotic systems through optimization of ribosome binding site sequences around the internal start codon^14,33^. At present, synthetic OLG expression in eukaryotic cells has not yet been demonstrated, though a number of strategies may be plausibly drawn from natural examples of multicistronic transcripts^34^. It is notable that OLGs in two of the three possible negative strand arrangements (-0 and -1) tend to score better than other frames. Opposite strand OLGs must be expressed by antisense transcription, which can counteract one another by a variety of mechanisms that can also be regulatory^35^. Thus, achieving efficient opposite strand OLG expression poses a unique problem but also an avenue for regulatory circuit design.

Lastly, our finding that natural protein domains can readily accommodate well defined folds without incurring significant or even any changes to their sequences suggests that novel OLGs are evolutionarily accessible via accumulation of mutations. This lends additional support to the emerging view of alternative reading frames as a potentially substantive contributor to *de novo* gene origin^36,37^. Altogether, the surprising ease with which we can design synthetic OLGs here leads us to conclude that at least in terms of the protein sequence space, the SGC does not severely limit the existence as well as evolution of OLGs in nature.

## METHODS

### Constrained iterative sampling algorithm for designing overlapping genes

We perform Gibbs sampling from generative models of the two given target proteins. One position is sampled per iteration for each protein, where the two positions are the residues translated by the overlapping triplet codons in the linked quartet representation of the nucleotide sequence. For each quartet, the pairwise product of the two amino acid probability vectors returned by the respective models gives the joint probability.

Depending on the frame arrangement, many pairs of amino acids are incompatible in the alternate reading frames and therefore their probabilities are hard constrained by zeroing them. The amino acid pair is sampled from this hard-constrained joint probability matrix, and the respective sequences are updated by replacing the current tokens with the sampled pair of tokens. In many cases, there are quartets that can encode the chosen pair of tokens, and all possible quartets are tracked. Because the quartets are linked by one nucleotide to its neighbors on both sides, some amino acids are incompatible with selected amino acids in neighboring positions. When the amino acid pair in the neighboring position(s) have already been chosen, the joint probability matrix is therefore additionally zeroed at the pairs that are incompatible with the neighboring residues. Choosing the amino acid pair for a position whose neighboring positions have already been chosen may also eliminate one or more of the possible quartets being tracked for the neighboring positions.

Each quartet / each position for the two target proteins are sampled exactly once. One sweep is when all positions have been visited; multiple sweeps may be performed. At the end of a sweep, pseudolikelihood (PLL) scores are obtained. PLL is defined by the summation of the conditional log probabilities of each token given all other tokens and is calculated by masking a position and predicting the probabilities for the masked position one at a time^38^. At the end of each sweep, the ratio of PLL scores are used to weigh the individual probability vectors in the next sweep before taking their pairwise product to balance the scores for the two proteins. Furthermore, the per-position conditional log probabilities calculated for PLL scoring are used to bias the order of positions to be sampled in the next sweep. This may be by the probabilities themselves such that higher mean (of the two amino acids across the positions) log probability positions are prioritized or by the entropies of the probabilities such that lower entropy positions are prioritized. We prioritize the scan order by entropy in all design runs shown. We perform 10 sweeps, look at the worst PLL scores of the pair of OLGs generated in each sweep and take the pair with the best of the worst PLL scores. The following pseudocode summarizes the algorithm:

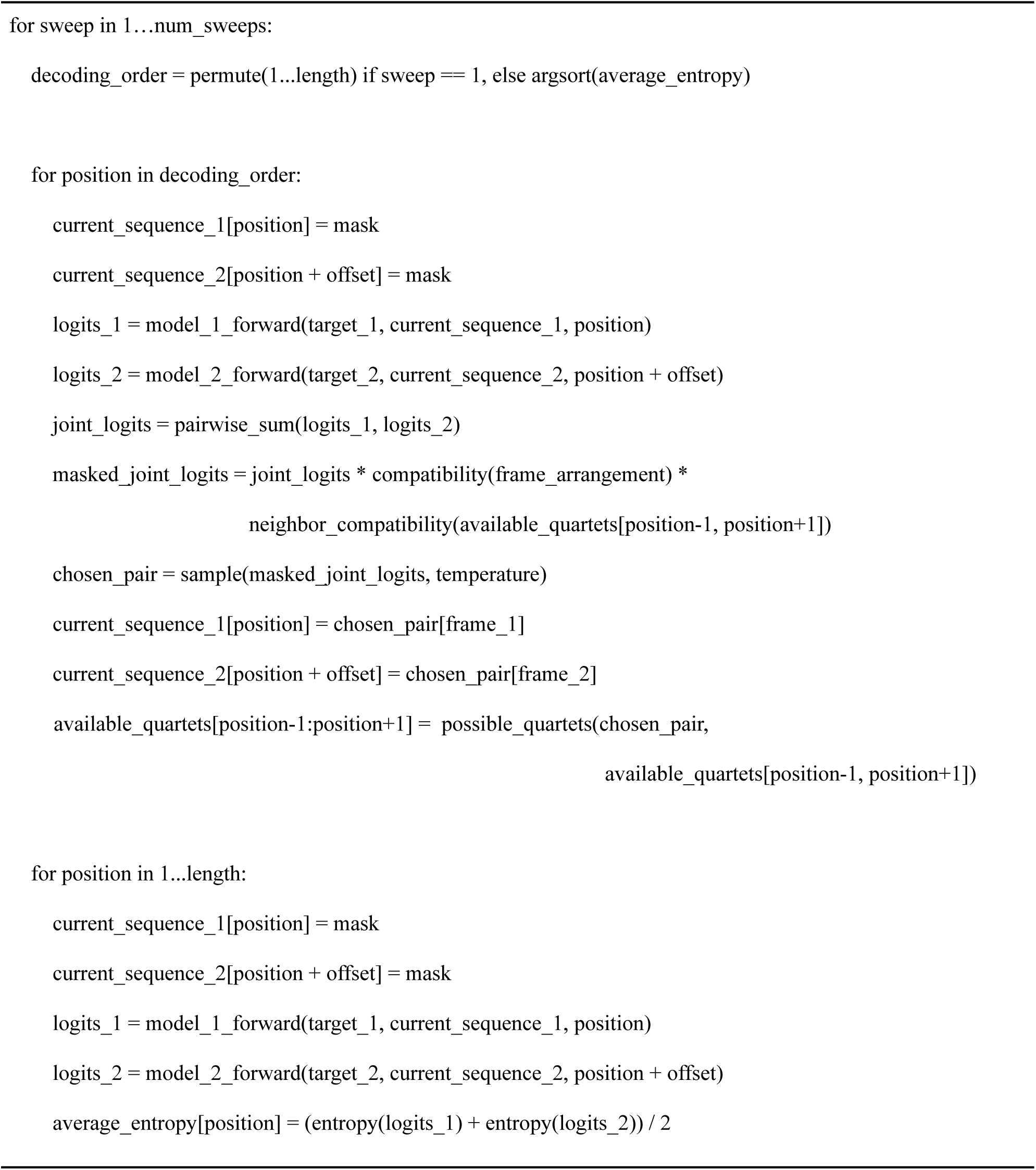

We observed frequent occurrences of repetitive amino acid spans when designing OLGs with some models even when using higher temperatures during sampling. To address this, we penalize repetitions by weighing down the log probabilities of the amino acids that were already selected beyond some threshold count within a specified window around a position. The weight is a penalty constant powered by the number of times the token has occurred within the window. We also employ a hard maximum percentage of the sequence that can be of single amino acid.

### Approximating the magnitude of overlap constraints

Analysis of the magnitude of overlap constraints in frames other than -0 are non-trivial since there is a phase shift which means that the constraints at any position are dependent on its neighboring positions. We can still however estimate the level of degeneracy by Monte Carlo approximation. We generate N=2000 random reference frame sequences, each for peptide lengths 1 to 8. We then enumerate all possible alternative frame amino acid sequences that are compatible with each random reference frame sequence.

To enumerate the possible sequences, we iterate through the linked quartet representation position-wise from left to right. At the first position, we list all possible alternative frame amino acids that are compatible with the fixed reference frame amino acid along with the possible last nucleotide(s) of the quartet encoding them. In the subsequent steps, we extend the subsequence by repeating this process for every possibility enumerated in the previous step. At the end of each step, the subsequences are deduplicated on the amino acid sequence and the last nucleotide of the last quartet. After the final step, the sequences are deduplicated only on the amino acid sequence, and the total number of unique possible sequences are counted.

We fit a linear model on the logarithm of the average number of possibilities versus peptide length. The base of the logarithm exponentiated by the coefficient of the fit gives the estimate of the conditional per-position degeneracy in the alternative frame.

### AlphaFold2 predictions

We run AF2 predictions using the AF2Rank protocol^25^. Specifically, we use single sequence inputs while providing a sidechain-masked template, zeroing the coordinates for sidechain atoms aside from Cβ and adding Cβ atoms to glycine residues. We used pTM model 2 or multimer model 2 (for CM1 homodimer predictions) and used a total of 6 recycles. We also run the single sequence predictions without templates using model 3 weights and take the prediction with the higher pLDDT. We use US-align for calculating TM scores and RMSD between target and predicted structures^39^. For CM1 / IF1 overlaps, template structures are taken from PDB accessions 1ECM and 3I4O.

### Designing CM1 / IF1 overlaps conditioned on multiple sequence alignments

Multiple sequence alignments (MSA) were obtained for CM1 and IF1 sequences (UniProt accessions P0A9J8 and P9WKK3) by ColabFold MMseqs2 query^40^. Sequences in the MSA with less than 75% coverage of the query sequence were dropped. We performed single sequence AF2 predictions with sidechain-masked template structures of the query proteins for the sequences in the MSA and further filtered them by average pLDDT threshold of 88. In each design run, we randomly subset 64 sequences in the filtered MSA to condition the generation, and the top row (query sequence) of the MSA is initialized with mask tokens at all positions. We use the EvoDiff-MSA model with MSA_OA_DM_MAXSUB weights. We use repetition penalty weight scaling factor of 1.1 with window size of 4 and cap the maximum proportion of the sequence that can be a single amino acid type at 25%. CM is slightly longer than IF1 (91 vs. 70 residues); we design all possible offsets which completely overlap the smaller IF1 within CM, prioritizing the scan order so that overlapping regions are sampled first in each sweep.

### Designing OLGs encoding the backbone structures

For the target proteins, we generated backbone structures using two different methods. First, we invert the RoseTTAFold (rf_Nov05_2021) structure prediction network by optimizing for input sequences that maximize the mean Kullback–Leibler divergence over the distance and angle distributions between the predicted structure and uniform background^26,41^. We also calculate and apply the radius of gyration term to the loss function, ranging from 13∼18Å, to vary the shapes of generated structures and favor a packed core^42^. We optimize the input first by gradient descent (GD) with softmax straight-through approximation and normalization of the gradient, using ADAM optimizer^43,44^. We then use the GD optimized sequences as seeds and iterate further by Markov Chain Monte Carlo sampling (MCMC). GD typically converges within 200 steps, after which we run up to 1000 MCMC steps.

We also generated monomeric backbones unconditionally via the RFdiffusion model, with the default parameters and fixing the length of the backbones to 100 amino acids^45^. We obtained a total of 1050 backbone structures with these two methods, annotated them by secondary structure class (α-only, β-only and mixed α-β) and randomly selected 5 structures from each class for 15 total. We then targeted all 15x15=225 possible pairwise combinations of these structures for OLG design with ProteinMPNN. We vary the offsets in between 0 to 5 which results in the overlap percentage of 95% at minimum. For fixing a specified proportion of the sequence in the reference frame, the positions to be fixed are chosen at random.

### Randomized genetic codes

For randomizing genetic codes, we use the strategies (amino acid permutation and All Amino Acids Get At Least One Codon shufflers) and the code from previous works^46–48^. We generate and use 1000 alternate randomized codes for each strategy. We use a smaller set of 3 target structures, 1 from each secondary structure class for 9 pairwise combinations; otherwise all parameters are the same.

### Pfam domains sequence set

We select a set of representative sequences from Pfam database version 37.0 by taking the first Pfam family for each clan and then taking the first sequence for each family’s seed alignment which is longer than 100 amino acids^49^. For each selected sequence, we randomly sample (length / 10) subregions which are 100 amino acids long. We use RandProt to make a Markov model from the final set and generate random protein sequences that preserve the k-mer distribution of the sequence set^50^.

### Protein language model embedding analysis

To obtain a vector representation of a protein (embedding), we extract the hidden states and average them across the positions in the sequence. We calculate the embeddings from ESM2 (esm2_t33_650M_UR50D), ESM3 (esm3-open-small) and ProstT5^21–23^. To project the embeddings into a two-dimensional space, we used the Uniform Manifold Approximation and Projection algorithm^51^. We use the following parameters: n_neighbors=15, n_components=2, metric=euclidean, n_epochs=200, init=spectral, min_dist=0.1.

### Cloning and recombinant expression

The 192 overlapping genes selected for experimental validation were synthesized as linear DNA fragments (eBlocks, Integrated DNA Technologies) and cloned into two plasmid vectors, each designed to express the protein coding sequence in either the +0 or +1 frame for a total of 384 unique proteins. These vectors are based on LM0627 (Addgene #191551) backbone^52^. They were modified to express the proteins in the different alternative reading frames while avoiding stop codons and minimizing charged or bulky amino acids that could interfere with the designed sequence (**Table S2**). Cloning, *E. coli* recombinant expression and purification were performed as previously described^45^. In short, each eBlock was cloned via Golden Gate assembly and transformed into chemically competent *E. coli* BL21 (DE3) cells. Four 1mL auto-induction cultures per construct were incubated at 37°C for 20 hours.

After incubation, cells were harvested and lysed. The clarified lysate, which serves to evaluate soluble cytoplasmic expression, was separated from the insoluble inclusion bodies. The clarified lysates were applied to Ni-NTA agarose resin in 96-well fritted plates and purified separately from the inclusion bodies. Inclusion bodies were solubilized by resuspending the insoluble fraction in 6M guanidine hydrochloride (GdnHCl) and refolding was done on beads by applying decreasing concentrations of GdnHCl (6M, 3M, 1.5M, 0.75M, 0.375M). Proteins were washed before elution and analyzed by size exclusion chromatography (SEC) on a Superdex75 Increase 5/150 GL column (Cytiva 29148722). Expression was considered successful if the protein yielded a clear peak at the expected molecular weight with a concentration above 2uM from a 4mL overnight E. coli culture. Proteins selected for further characterization were purified at a larger scale from 50mL cultures and analyzed by SEC on a Superdex75 Increase 10/300 GL column (Cytiva 29148721) as previously described^52^. To assess secondary structure and thermostability, purified proteins were analyzed by circular dichroism (CD) as previously described with scans from 190 nm to 260 nm at 10°C intervals between 25°C and 95°C^53^.

## Acknowledgements

This research was supported by an appointment to the Intelligence Community Postdoctoral Research Fellowship Program at the University of Washington administered by Oak Ridge Institute for Science and Education through an interagency agreement between the U.S. Department of Energy and the Office of the Director of National Intelligence (GB). ME acknowledges support from la Caixa Foundation and Rafael del Pino Foundation. DB acknowledges support from the Audacious Project at the Institute for Protein Design (PG117878: Audacious Hub); the National Institutes of Health’s (NIH) National Cancer Institute grant R01CA114536 (GR024635: FHCC RIDDELL R01 - 66-8725 - 2021), and R01CA240339 (GR009231: NCI BAKER NKC R01 - 62-0055 - 2021). GS acknowledges support from NIH Awards R33CA286947, R56HG013312 and R01GM149631.

## Author contributions

Conceptualization: GB, GS; Methodology: GB, ME; Investigation: GB, ME; Visualization: GB, ME; Funding acquisition: GB, GS; Project administration: GB; Supervision: DB, GS; Writing - original draft: GB, ME; Writing - review & editing: GB, ME, DB, GS

## Competing interests

Authors declare that they have no competing interests.

## Code availability

The code implementing the algorithm described in this work is available at this Github repository: https://github.com/gwbyeon/olgdesign.

## Supplementary FIGURES

**Supplementary Figure 1.**
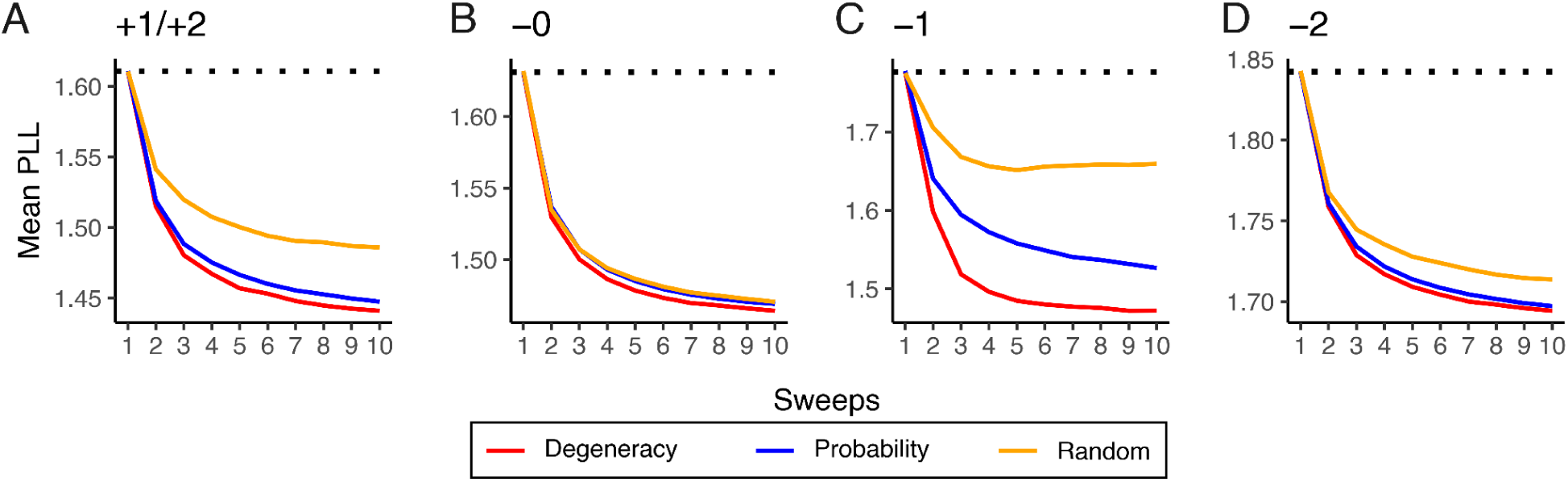
Computational design of synthetic overlapping genes. (**A**) The effects of biased scan orders on pseudolikelihood scores of the designed sequences. Pseudolikelihoods of the designed sequences are plotted along multiple complete sweeps through the positions. Colors indicate different strategies to determine the scan order of the positions in the successive sweeps. (**B**)-(**D**) same for other frames.

**Supplementary Figure 2.**
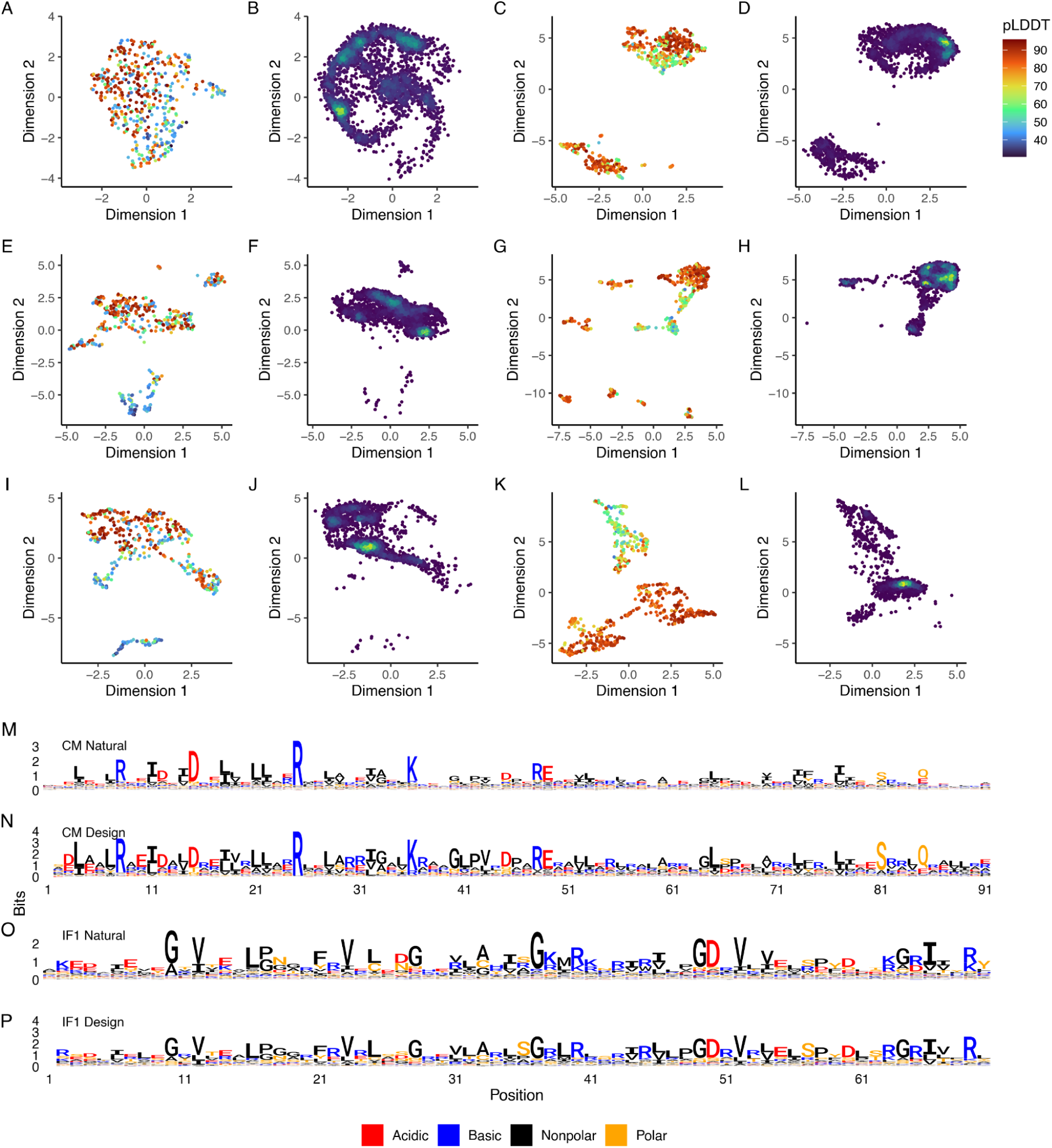
Targeting a pair of homologous protein families for OLG sequence design. (**A**) UMAP visualization of ProstT5 embeddings of the natural homologous sequences of CM. Colored by mean pLDDTs of the AlphaFold2 predictions for each sequence. (**B**) Projection of the ProstT5 embeddings of the OLG synthetic homologs of CM in the same space of (A), colored by density. (**C**, **D**) Same as (A) and (B) but for IF1 sequences. (**E**-**H**) Same as (A)-(D) but with ESM3 embeddings. (**I**) A sequence logo of the position weight matrix derived from the multiple sequence alignment of the natural homologues of CM. The height of each letter is scaled proportionally to entropy. Colored by acidic, basic, polar or nonpolar side chain types. (**J**) Sams as (I) but for the designed overlapping sequences for CM. (**K**, **L**) Same as (I) and (J) but for IF1.

**Supplementary Figure 3.**
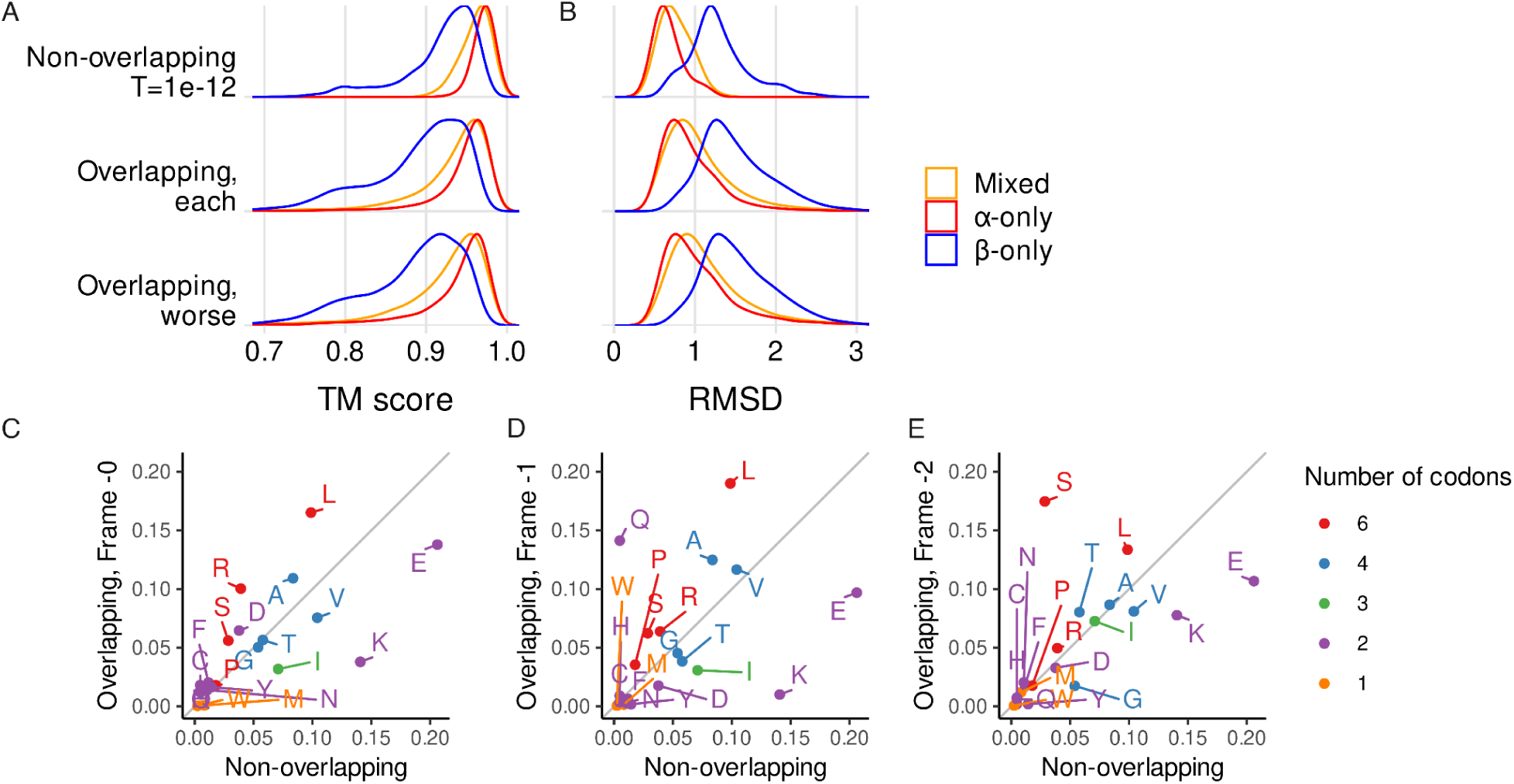
Designing OLG sequences encoding pairs of highly ordered protein structures. (A) Distribution of TM scores for the AlphaFold2 predictions of ProteinMPNN designed sequences versus the target backbone structures, as overlapping pairs or standard non-overlapping proteins. Overlapping designs are shown altogether (middle), or only the worse TM scores of the overlapping pair are plotted (bottom). Colored by secondary structure classes. (B) Same as (A) but plotting the distribution of RMSDs. (**C**) A scatter plot of the amino acid compositions of the designed overlapping (frame -0) versus non-overlapping sequences. Each point labeled with amino acid letters and colored by the number of codons for the amino acid in the standard genetic code. (**D**, **E**) Same as (C) but for frames -1 and -2, respectively.

**Supplementary Figure 4.**
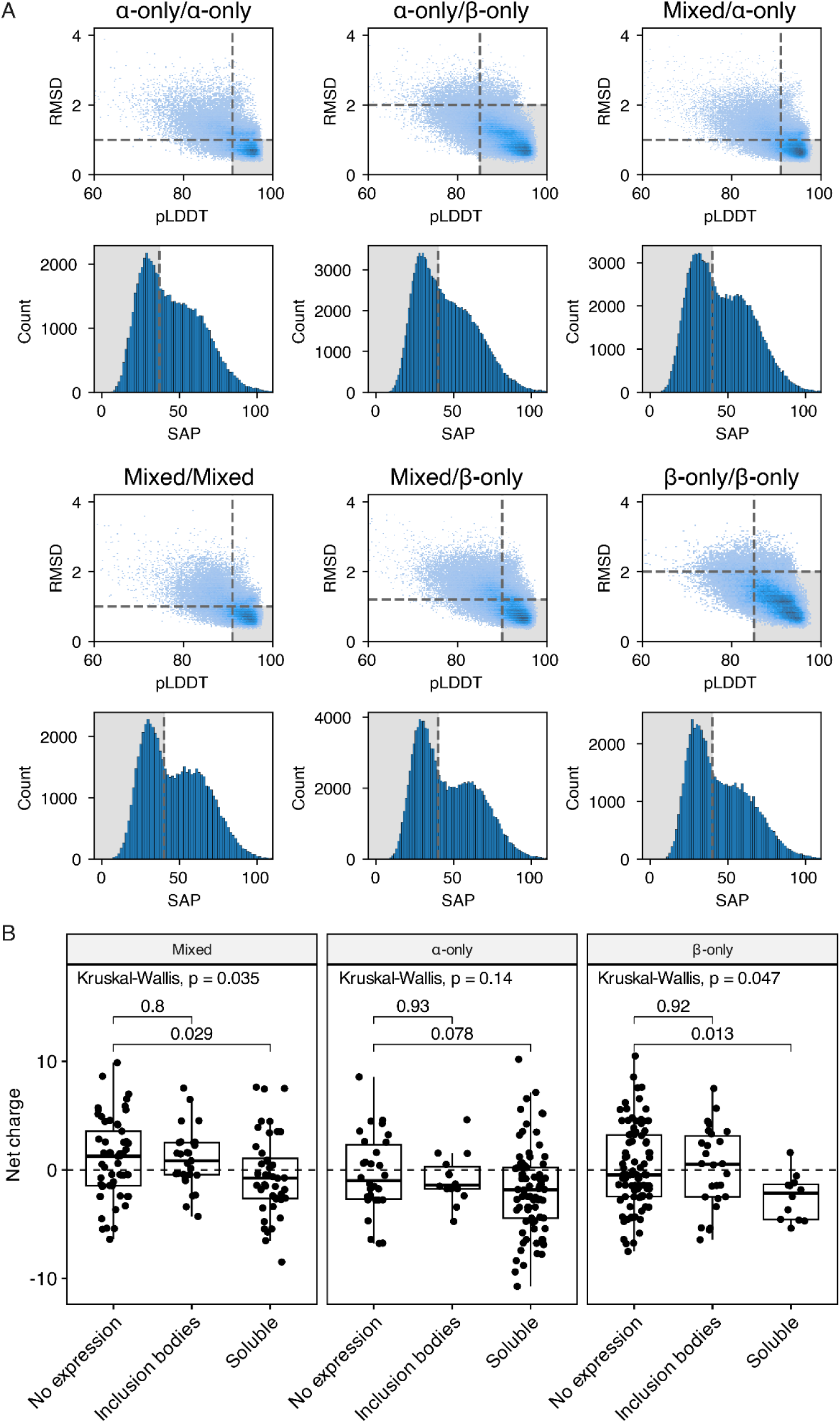
Experimental characterization of overlapping *de novo* protein pairs. (A) *In silico* metrics by secondary structure class and thresholds applied for filtering. The 32 selected designs for each category were randomly sampled from the intersection of the shadowed areas in both plots. (**B**) Net charge of the proteins by recombinant expression success and secondary structure class. The lower and upper hinges correspond to the first and third quartiles. The midline corresponds to the median. The whiskers are bounded by 1.5 IQR. Labeled numbers show two-sided Wilcoxon p-value between groups indicated by the line or Kruskal-Wallis p-value at the top.

## SUPPLEMENTAL TABLES

**Table S1.** Thresholds for filtered designs.

| | α-only/<br>α-only | α-only/<br>β-only | Mixed/<br>α-only | Mixed/<br>mixed | Mixed/<br>β-only | β-only/<br>β-only |
| --- | --- | --- | --- | --- | --- | --- |
| pLDDT | >91 | >85 | >91 | >91 | >90 | >85 |
| RMSD | <1 | <2 | <1 | <1 | <1.2 | <2 |
| SAP | <37 | <40 | <40 | <40 | <40 | <40 |
| # passing | 51 | 1312 | 442 | 132 | 331 | 136 |

**Table S2.** Vectors for overlapping gene expression. Key differences between the vectors are highlighted. The four residue overhangs during Golden Gate cloning are in uppercase. The 3’ adapter used for each eBlock depends on the last base of the overlapping gene insert, affecting the cloning product and amino acids adjacent to the C-terminus.

| Vector | Cloning product | Expression product |
| --- | --- | --- |
| +0 frame | insert ending in G/A:<br>...atgAGGA<u>tc</u>-insert(g/a)-**g**TTCCggc...<br>insert ending in T/C:<br>...atgAGGA<u>tc</u>-insert(t/c)-**c**TTCCggc... | insert ending in G/A:<br>**MRI**-insert-(**G/S**)SG...<br>insert ending in T/C:<br>**MRI**-insert-(**S/P**)SG... |
| +1 frame | insert ending in G/A:<br>...atg<u>tc</u>AGGA-insert(g/a)-**g**TTCC<u>tg</u>gc...<br>insert ending in T/C:<br>...atg<u>tc</u>AGGA-insert(t/c)-**c**TTCC<u>tg</u>gc... | insert ending in G/A:<br>MSG-insert-**V**PG...<br>insert ending in T/C:<br>MSG-insert-**L**PG... |

## Notes

### Competing Interest Statement

The authors have declared no competing interest.

